# Live Imaging of Cutaneous Wound Healing in Zebrafish

**DOI:** 10.1101/2022.11.07.515499

**Authors:** Leah J. Greenspan, Keith Ameyaw, Daniel Castranova, Caleb A. Mertus, Brant M. Weinstein

## Abstract

Cutaneous wounds are common afflictions that follow a stereotypical healing process involving hemostasis, inflammation, proliferation, and remodeling phases. In the elderly or those suffering from vascular or metabolic diseases, poor healing following cutaneous injuries can lead to open chronic wounds susceptible to infection. The discovery of new therapeutic strategies to improve this defective wound healing requires a better understanding of the cellular behaviors and molecular mechanisms that drive the different phases of wound healing and how these are altered with age or disease. The zebrafish provides an ideal model for visualization and experimental manipulation of the cellular and molecular events during wound healing in the context of an intact, living animal. To facilitate studies of cutaneous wound healing in the zebrafish, we have developed an inexpensive, simple, and effective method for generating reproducible cutaneous injuries in adult zebrafish using a rotary tool. Using our injury system in combination with live imaging, we can monitor skin re-epithelialization, immune cell recruitment, and vessel regrowth and remodeling in the same animal over time. This injury system provides a valuable new experimental platform to study key cellular and molecular events during wound healing *in vivo* with unprecedented resolution.

## Introduction

Cutaneous wound healing is a complex process that requires the coordination of many different cell types to ensure wound closure and tissue repair. It is characterized by four overlapping phases: hemostasis in which a blood clot is formed, inflammation where immune cells are recruited to the site of injury, proliferation which consists of skin re-epithelialization, angiogenesis, and fibrogenesis, and remodeling in which vessels are pruned and collagen remodeled, usually resulting in the formation of a scar (Schrementi et al., 2015). Delayed and/or ineffective healing after cutaneous injuries is common in the elderly and those suffering from vascular disease or diabetes, and it can lead to chronic wounds susceptible to infection (Guo and Dipietro, 2010, Schrementi et al., 2015). Better understanding the cellular and molecular dynamics of the different wound healing phases and how they are altered during aging or disease conditions is critical to generate better therapeutics to ameliorate defective healing.

Mammalian models of wound healing have revealed many important insights into the cell types and signals that drive the different phases of wound healing. However, challenges in carrying out continuous, high resolution live imaging and experimental manipulation of the healing process in living, intact mammalian models have limited the usefulness of these models. Zebrafish provide an attractive vertebrate model for studying tissue injury and repair. They have a high regenerative capacity in many different tissues and organs, including the heart and caudal fin (Johnson and Weston, 1995, Poss et al., 2002), and major stages of cutaneous wound healing are conserved in zebrafish with the added advantage that zebrafish tend to have minimal scar formation (Richardson et al., 2013). Unlike mammals, zebrafish have optically clear embryos and larvae, and pigment-free adult zebrafish lines are available (White et al., 2008), facilitating high-resolution optical imaging of cellular and even subcellular features in intact animals. We have also developed methods for long-term time-lapse imaging of intubated adult fish (Castranova et al., 2022) that permit extended visualization of cellular and subcellular dynamics in living, healthy adult animals. The intrinsic imaging potential of the fish is complemented by the availability of a plethora of transgenic and mutant fish lines that permit direct visualization and functional manipulation of many, if not all, of the different epithelial, immune, vascular, and other cell populations that contribute to the wound healing process. Together these features make zebrafish a valuable vertebrate model for direct observation of the dynamics of wound healing and for uncovering the cellular and molecular processes driving tissue repair after cutaneous injury.

Previous attempts to study cutaneous injury in the zebrafish have involved using either a specialized, expensive dermatology laser to create wounds (Richardson et al., 2016, Richardson et al., 2013), or fine forceps to create small and inconsistent superficial laceration or abrasion wounds (Noishiki et al., 2019, Yuge et al., 2022). Here, we report an inexpensive, simple, and effective method for generating reproducible cutaneous abrasion injuries in adult zebrafish using a rotary tool. We describe and provide detailed Computer Aided Designs (CAD) for custom 3D printed items that allow for better reproducibility such as a cowling to control injury depth, and a fish holding platform with a rotating insert that facilitates immobilization and positioning for better control of injury placement. We demonstrate the usefulness of this method for studying the wound healing process by generating cutaneous injuries in transgenic zebrafish and then using high-resolution live imaging to monitor skin re-epithelialization, neutrophil recruitment, and blood and lymphatic vessel regrowth into the wound site. By performing repeated imaging of the same fish over a two-week span after injury we show that it is possible to observe in detail and accurately assess the timing of wound healing phases. Our method will facilitate imaging and experimental analysis of cutaneous wound healing in zebrafish, and can be used to study changes in wound healing in aged fish or in readily available diabetic fish models.

## Results

We developed an easy to use, inexpensive and reproducible abrasion injury method for adult zebrafish using a rotary tool (**Fig. 1A**). The rotary tool is mounted on a workstation comprised of a drill press, rotary tool holder, and mounting stand, allowing the rotary tool to be mounted at an angle and smoothly and incrementally advanced towards the sample. A square nose 1/16” end mill cutting bit (**Fig. 1B,C**) is mounted on the end of the tool, and a custom 3D printed plastic cowling (**Fig. 1D, Supp. File 1**) is then attached to the tip of the rotary tool (**Fig. 1E**) around the bit, with only a tiny amount of the tip of the cutting bit protruding from the cowling. This custom printed cowling helps limit and control the depth of the abrasion injury (see methods). The entire injury apparatus is placed next to a dissecting microscope. Fish are mounted in a custom 3D printed plastic holding platform with a rotating plastic insert (**Fig. 1F, Supp. File 2**) that allows the angle of the animal to be adjusted for better control of injury placement. During injury the holding platform is placed on top of the dissecting microscope base, adjacent to the rotary tool, as shown (**Fig. 1G**).

**Figure 1:**
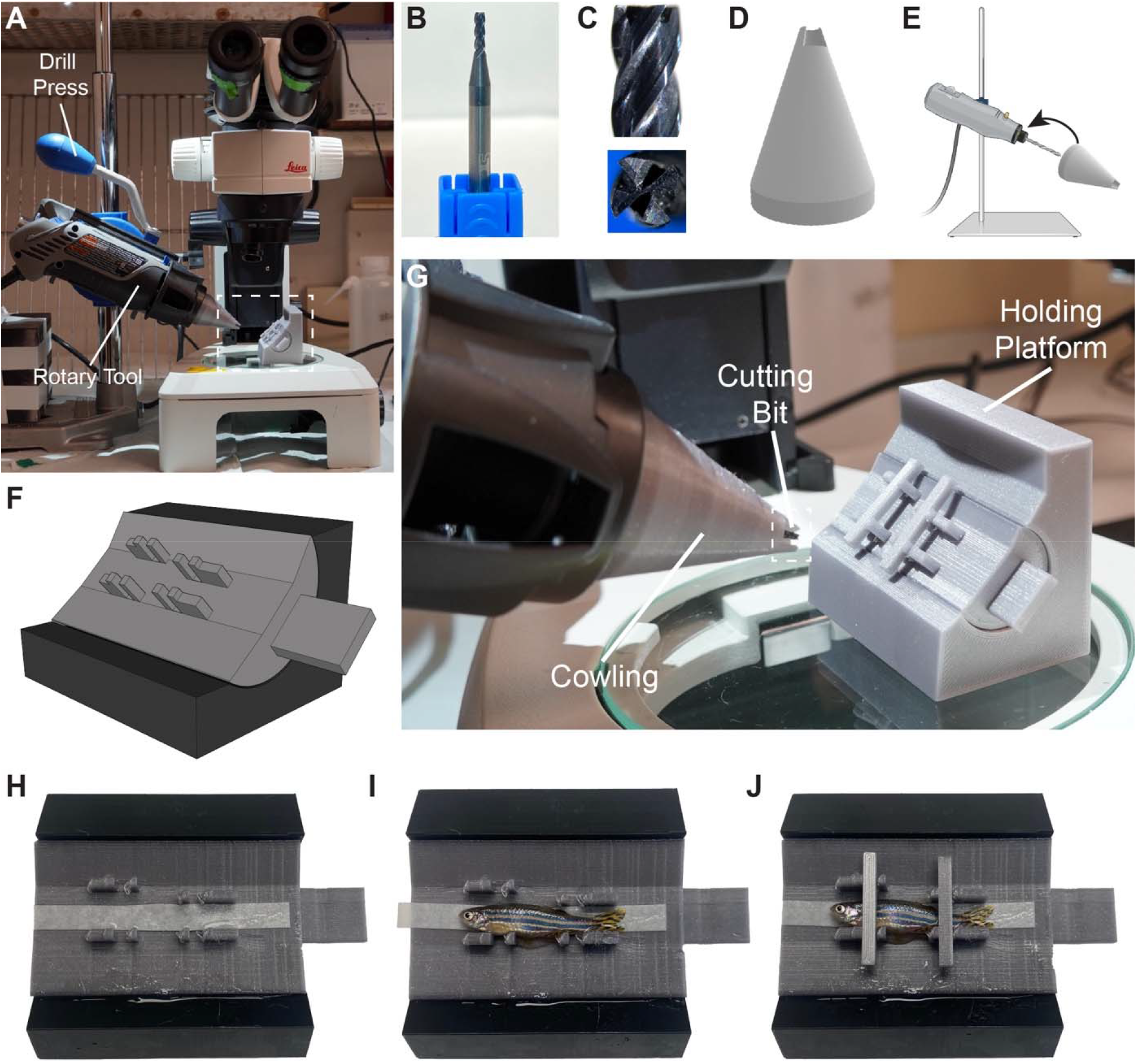
Cutaneous wounding using a rotary tool. **A.** Overview image of the apparatus used for generating cutaneous abrasion injuries. **B.** 1/16^th^ inch square end mill cutting bit used for generating cutaneous wounds. **C.** Higher magnification side and front view of the tip of the cutting bit. **D.** 3D rendering of the plastic printed cowling used to control injury depth. **E.** Schematic diagram showing attachment of the cowling to the rotary tool. **F.** 3D rendering of the plastic printed holding platform with rotating insert used to anchor the fish during injury. **G.** Higher magnification image of the rotary tool with cutting bit, cowling, and holding platform on a dissecting microscope. **H-J.** Images of mounting of a fish onto the holding platform, showing a platform with wetted filter paper (H), a fish placed on the platform on top of the paper (I), and a mounted fish immobilized on the platform with anchoring rods (J).

Adult fish being subjected to abrasion injury are anesthetized in system water containing 0.5X Tricaine for 4 minutes, then transferred to system water containing 1X Tricaine for up to a minute longer, until they cease to move. The fish are then removed using a plastic spoon and carefully mounted on a holding platform containing a strip of filter paper soaked in system water containing 1X Tricaine (**Fig. 1H-J, Movie 1**). Fish are placed on the platform on top of the wetted paper (**Fig. 1I, Movie 1**), and then immobilized in place using custom 3D-printed anchoring rods (**Supp. File 3-4**) that fit securely between pegs designed into the holding platform (**Fig. 1J, Movie 1**). The rotatable insert the fish is mounted on in the holding platform allows the angle of the fish to be adjusted for better control of placement of the injury. Once the fish is fully mounted, the holding platform is placed on the stereoscope and the rotary tool positioned to ensure proper placement of the injury site on the left flank of the fish between the pelvic and anal fins. The rotary tool is turned on at speed 3 (8500 RPM) and pressed gently into the side of the fish for ten seconds, creating a 1/16” diameter abrasion injury approximately 2.4 × 10^6^ μm^2^ in area (**Fig. 2A,B, Movie 1**). After injury, fish are quickly removed from the holding platform by pulling forward on the filter paper and releasing the fish into a tank of fresh system water where water is moved over the fish until it has awakened. 16/16 fish that we subjected to this procedure quickly recovered from anesthesia after injury and were swimming normally shortly after revival. They displayed no obvious signs of distress, and most (13/16) of the fish were still alive and appeared healthy 2 weeks after injury. Those fish that died, were due to repeat Tricaine exposure from imaging and not due to the injury itself.

**Figure 2:**
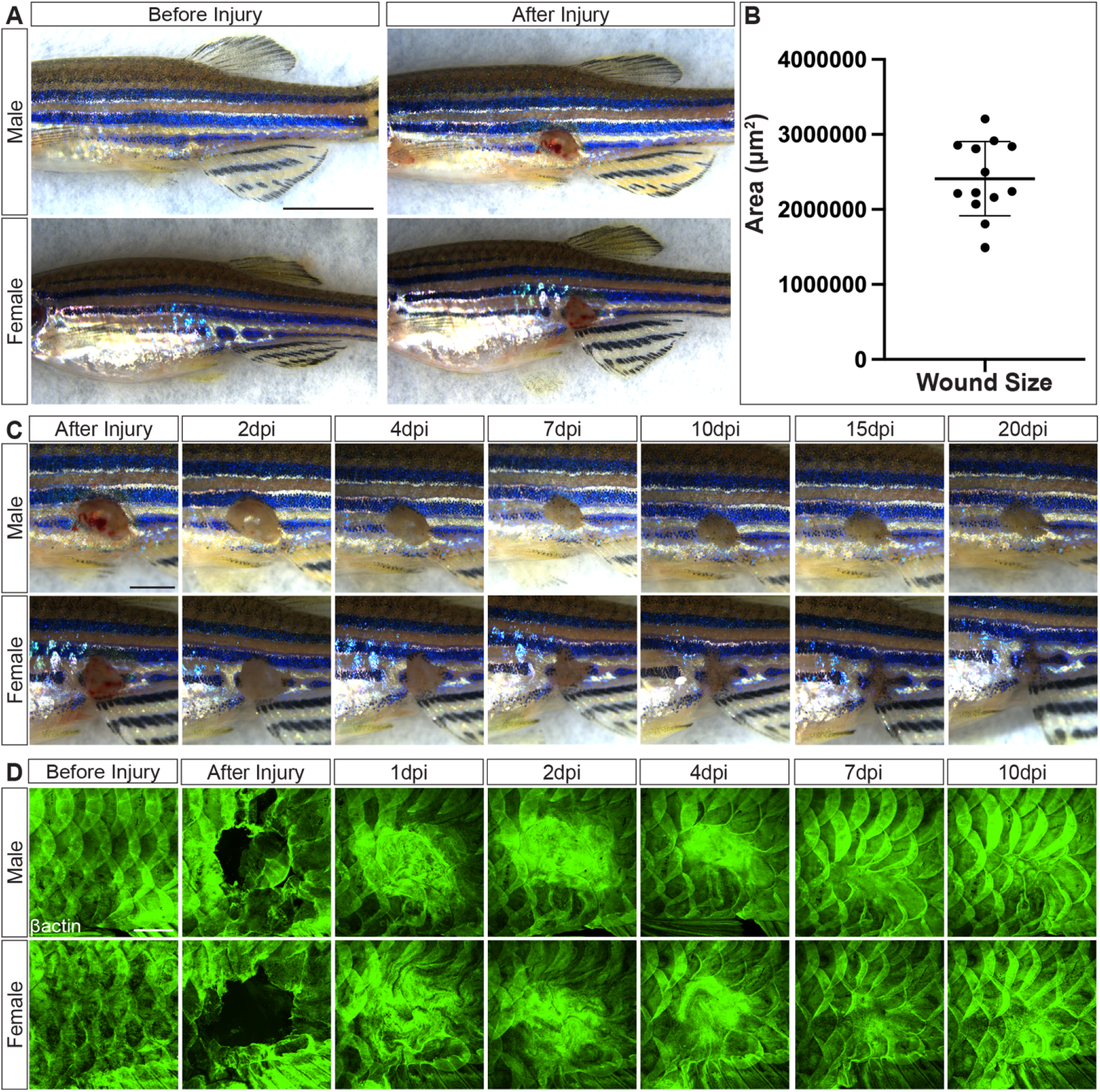
Reproducible cutaneous wounds can be made with a rotary tool. **A.** Images of a male and female fish before and after cutaneous wounding, showing consistent wound formation using a rotary tool. **B.** Quantification of wound area, with each dot representing a single fish (n=13 fish). Error bar indicates mean area with standard deviation. **C.** Magnified images of the flanks of male (top) and female (bottom) immediately before and after cutaneous injury and at intervals thereafter up to 20 days post injury. **D.** Confocal images of the flanks of *Tg(actb2:GFP)* (βactin) transgenic male (top) and female (bottom) fish immediately before and after cutaneous injury and at intervals thereafter up to 10 days post injury. All images are oriented rostral to the left and dorsal up. Scales bars depict 5000μm (A), 2000μm (C), and 1000μm (D).

We collected a time series of stereoscope images of the male and female fish shown in Fig. 2A-D, collecting images for 20 days after injury **(Fig. 2C)**. Blood clotting occurs rapidly, with cessation of bleeding almost immediately after wound induction (**Fig. 2C**, after injury). As previously reported for zebrafish (Richardson et al., 2016, Richardson et al., 2013), wounds were re-epithelialized by 1 day post injury (dpi), and the unpigmented area continued to grow smaller throughout the course of the 20 day time series (**Fig. 2C**, 2 dpi to 20 dpi). To examine re-epithelialization directly and with higher resolution, we collected a confocal time series of injured *Tg(actb2:GFP)* transgenic adult animals with β-actin expressing cells in green **(Fig. 2D)**. Prior to injury, GFP expression is mostly in the scales (**Fig. 2D**, Before Injury), with loss of GFP in the wound area after injury induction (**Fig. 2D**, After Injury). By 1 dpi, GFP is re-expressed throughout the wound site suggesting that the wound has been sealed (**Fig. 2D**, 1 dpi). Timelapse confocal imaging of an injured *Tg(actb2:GFP)* transgenic adult fish from 0.5-17.5 hours post injury (hpi) shows a stream of epithelial cells moving into the wound area directly after injury, confirming rapid wound closure (**Movie 2**). This agrees with previous reports showing that unlike in mammals, where re-epithelialization occurs in later phases of wound healing, zebrafish re-epithelialize their wounds as one of the first steps of the repair process (Richardson et al 2013, Richardson et al 2016). After initial wound closure, scales can be seen gradually regrowing over the injury site, covering the entire wound by 10 dpi (**Fig. C,D**, 1 dpi to 10 dpi).

In addition to epithelial cells, high-resolution confocal imaging of injured *Tg(actb2:GFP)* fish suggested other, highly motile cell types were also infiltrating healing wounds (**Movie 2**). To examine whether some of these might be neutrophils recruited to the injury site, we collected high-resolution confocal images of a *Tg(lyz:dsred)* transgenic, *casper* mutant (pigment-deficient) adult fish, as a continuous time-lapse from 0.5-17.25 hpi **(Movie 3)**, and then at intervals thereafter until 15 days post-injury **(Fig. 3)**. Although neutrophils were present on the surface of the fish prior to injury **(Fig. 3A,I)**, there was a rapid influx of a large number of neutrophils into the wound site after injury **(Movie 3, Fig. 3C,K)**. There was already extensive migration of neutrophils to the periphery of the wound by 2 hours post-injury **(Movie 3, Fig. 3B,J,Q)**, with neutrophil recruitment peaking by 1 dpi **(Fig. 3C,K,Q)**. This initial burst of neutrophil recruitment was followed by a more gradual decline in neutrophil localization to the wound site **(Fig. 3D-Q)**, with neutrophil levels close to baseline by 15 dpi **(Fig. 3H,I)**. This suggests that zebrafish have an acute neutrophil response to cutaneous abrasion wounds that dissipates quickly.

**Figure 3:**
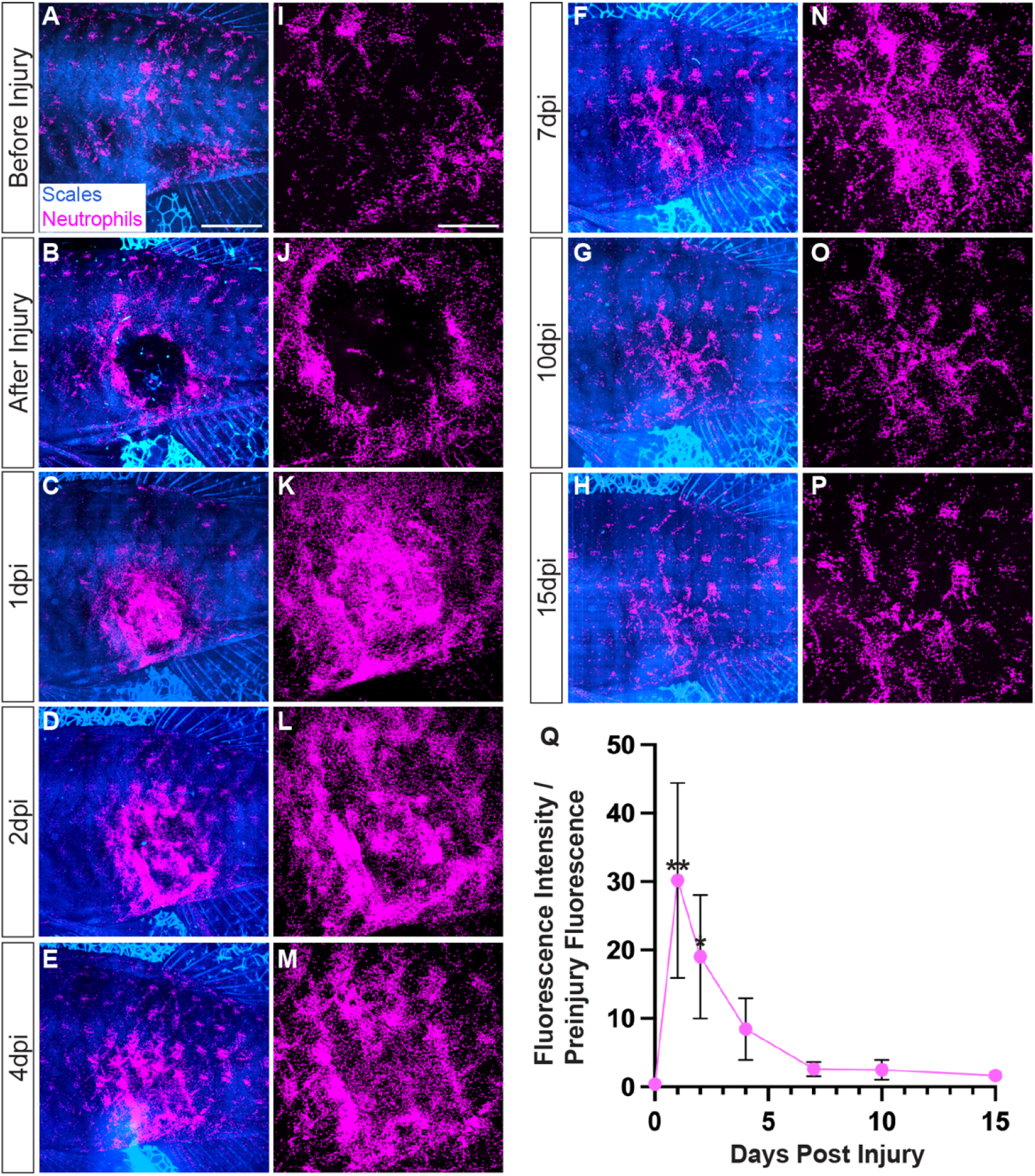
Neutrophils are rapidly but transiently recruited to cutaneous wounds. **A-P.** Maximum projection confocal images of adult *Tg(lyz:DsRed2)^nz50^* transgenic *casper* mutant fish with scales in blue (autofluorescence) and neutrophils in magenta (lyz:dsRed). Panels I-P show magnified views of lyz:dsRed-positive neutrophils in the wound site. I. Quantification of the mean fluorescence intensity of lyz:dsRed-positive neutrophils in the wound site, normalized to preinjury neutrophil fluorescence. Each dot represents the mean ratio of 4 fish at the indicated time point. A strong increase in neutrophil density measured by fluorescence intensity is seen by 1 day post injury, but it declines gradually thereafter. Error bars indicate mean ratio and standard error. Ratio paired t test, **p<0.01, *p<0.05. Scale bars depict 2000μm (A-H) and 1000μm (I-P).

Angiogenesis is a key part of the wound healing process, ensuring proper reperfusion and restoration of blood flow to healing tissues and drainage of accumulated fluid. We examined blood and lymphatic revascularization following cutaneous injury in *Tg(mrc1a:egfp), Tg(kdrl:mcherry)* double-transgenic, *casper* mutant adult fish over the first 15 days post-injury. In uninjured adult zebrafish superficial lymphatic vessels form a stereotypic pattern, with the lateral lymphatic vessel running along the length of the fish at the midline horizontal myoseptum, and superficial intersegmental lymphatic vessels extending dorsal and ventrally away from the lateral lymphatic vessel along each of the myotomal boundaries (**Fig. 4A-B,E,K**). Skin blood vessels have a more irregular and tortuous pattern, with some thicker branched bundles, although underlying muscle blood vessels are thinner and align longitudinally across myotomes (**Fig. 4A,C,E,Q**). Vessel growth into wounds is rapid, with greater than 75% of wound area revascularized by 4 dpi (**Fig. 4D,E-H,K-N,Q-T**). Immediately after injury wounds contain only some severed fragments of blood and lymphatic vessels (**Fig. 4F,L,R**), some of which disappear by 2 dpi (**Fig. 4G,M,S**). Sprouting angiogenesis initiates by 2 dpi, continuing at a fast rate until 4 dpi when the regrowth rate begins to slow as vessels continue to infiltrate the wound area (**Fig. 4D,G-H,M-N,S-T**). Vessel growth continues at a slower pace for several more days, with most wounds having approximately 90% blood and lymphatic vessel coverage by 10 dpi (**Fig. 4I-J,O-P,U-V**), although remodeling of vessels continues for several more months (data not shown). Together this data shows that cutaneous wounds are efficiently revascularized after abrasion injuries using a rotary tool, with a rapid initial blood and lymphatic vessel outgrowth in the first few days followed by slower phases of growth and remodeling.

**Figure 4:**
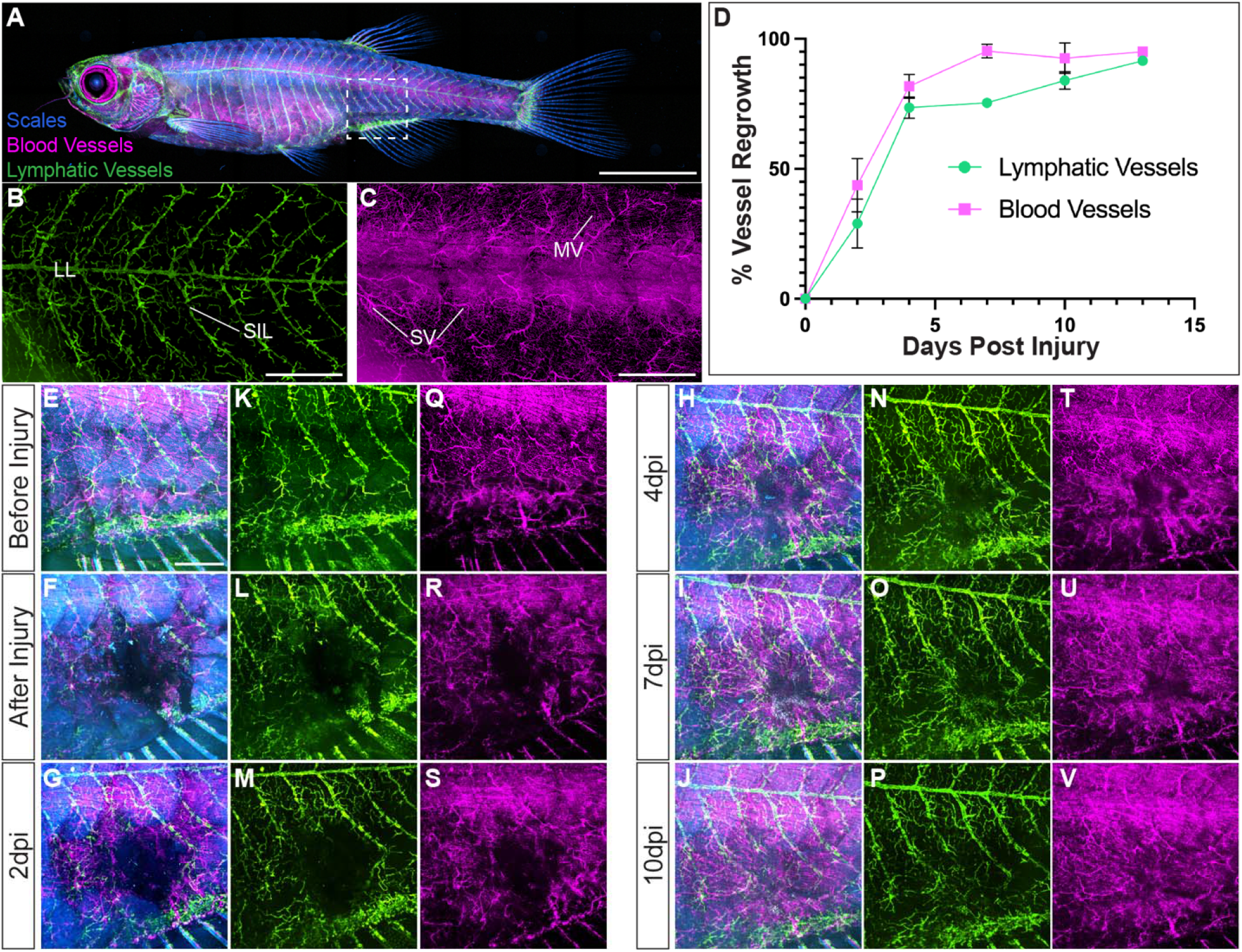
Revascularization of cutaneous wounds. **A-C.** Confocal images of an uninjured *Tg(mrc1a: eGFP)^y251^, Tg(kdrl:mcherry)^y206^* double transgenic, *casper* mutant adult fish with scales in blue (autofluorescence), lymphatic vessels in green (mrc1a:eGFP), and blood vessels in magenta (kdrl:mCherry). Whole-fish overview (A) and magnified flank (B-C) confocal images are shown, with B and C showing only lymphatic (green) or blood (magenta) vessels, respectively. White dotted box in A indicates approximate site of injury for E-V. **D.** Quantification of the % of wound area revascularized by blood (magenta) or lymphatic (green) vessels at time points following cutaneous injury (n=4 fish). Error bars indicate mean % and standard errors. **E-V.** Magnified confocal images of the trunk of the same *Tg(mrc1a: eGFP)^y251^, Tg(kdrl:mcherry)^y206^* double-transgenic *casper* mutant adult fish before cutaneous injury (E, K, Q), immediately after injury (F,L,R), and at several time points thereafter through 10 dpi (G-V). E-J show scales (blue autofluorescence), lymphatic vessels (green mrc1a:eGFP-positive), and blood vessels (magenta kdrl:mCherry-positive), while K-P and Q-V show only lymphatic vessels (green) or blood vessels (magenta), respectively. Scale bars depict 5000μm (A), and 1000μm (B-C, E-V).

## Discussion

We have developed a new method to generate cutaneous wounds in adult zebrafish. Using a rotary tool mounted on a drill press, along with custom 3D printed items including a cowling which attaches to the rotary tool to control the depth of injury and an adjustable holding platform to immobilize fish for the procedure, we can create reproducible cutaneous abrasion injuries with an easy to use, inexpensive setup. We have included 3D CAD files for custom 3D printing of the cowling, holding platform, and anchoring rods to allow other labs to easily reproduce our methods (**Supp Files 1-4**). Using our cutaneous abrasion injury method, we can easily monitor the initiation, progress, and duration of different phases of wound healing by either continuous time-lapse imaging of intubated adult fish (Castranova et al., 2022) or serial imaging of the same fish at multiple time points after injury. As has been noted using other injury methods in zebrafish (Richardson et al., 2016, Richardson et al., 2013), re-epithelialization of rotary tool induced abrasion wounds and recruitment of neutrophils occurs very rapidly, within a few hours after injury, suggesting this method generates wounds that efficiently trigger proper wound healing responses. Re-epithelialization and immune cell recruitment are followed by a revascularization phase that also occurs quickly over the first four days post-injury, with slower growth thereafter transitioning into an extended phase of vascular remodeling. Together, these findings suggest that our new cutaneous abrasion injury model combined with sophisticated methods for live imaging of transgenic adult zebrafish will yield important new insights into changes in cell behavior that occur during the wound healing process.

Previous reports suggest that zebrafish undergo many of the same stages of wound healing found in mammals (Richardson et al., 2013). However, mammals are generally euthanized for high-resolution analysis of healing tissues using immunofluorescence, immunohistochemistry, or other non-vital methods. In the zebrafish thousands of transgenic lines are available for an enormous variety of different cell types including epithelial cells, immune cells, and blood and lymphatic vessels, facilitating high-resolution live imaging of the healing process over time in the same animal (Choe et al., 2021, Gore et al., 2011, Hall et al., 2007, Jung et al., 2017). Combinations of transgenic lines labeling different cell types can be utilized to understand cellcell interactions occurring during the wound healing process. Molecular reporter lines have also been generated in zebrafish for Bmp, TGFβ, Notch, Shh, Erk, and other signaling pathways, permitting real-time assessment of these pathways during the wound healing process (Okuda et al., 2021, Schiavone et al., 2014). Together, these tools make the zebrafish a powerful model to study the coordination and dynamic nature of the cellular and molecular processes driving cutaneous wound healing.

Delayed wound healing can lead to the open chronic wounds susceptible to infection common in aging individuals and those suffering from vascular disease or metabolic disorders such as diabetes (Guo and Dipietro, 2010, Schrementi et al., 2015). Zebrafish exhibit signs of aging similar to those in mammals such as bone and muscle degeneration, cardiovascular impairment, and cognitive decline (Keller and Keller, 2018). Several highly effective zebrafish models are also available for modeling mammalian diabetes (Gleeson et al., 2007, Moss et al., 2009). These models and the transgenic lines available in zebrafish provide powerful tools for studying the effects of age and metabolic disorders on cutaneous wound healing. Combined with these tools, our new method for generating cutaneous wounds in zebrafish could have wide ranging implications for understanding the cutaneous wound healing process, facilitating the formation of new and improved therapies.

## Materials and Methods

### Wound healing apparatus

CAD designs are provided for custom 3D-printed plastic holding platforms and mounting rods used to hold male or female adult zebrafish in place for cutaneous injury, and for the cowling used to control the injury depth (**Supp. Files 1-4**). The holding platform consists of an outer holder with a 3D printed rotating insert the fish is mounted on (**Supp. File 2**). A small amount of silicone grease (SG-ONE 24708) was applied to the slot in the base before the rotating insert is inserted. The grease allows the rotating insert to move smoothly, and also prevents movement of the insert during injury. Printed anchoring rods are used to keep the fish tightly secured (**Supp. File 3-4**). The rotating insert allows for easy positioning of the fish for the injury procedure. The 3D-printed cowling is designed to attach to the end of the rotary tool with only a very small part of the bit tip exposed to limit the depth of injuries (**Supp. File 1**). 3D models of the holding platform, anchoring rods, and cowling were designed using SketchUp 3D modeling software and printed out of polylactic acid (PLA) using an Original Prusa i3 MK3S+ 3D printer (Prusa Research). These CAD designs could also be printed by commercial suppliers such as Xometry (xometry.com). The rotary tool was a Dremel 400 XPR attached to a Dremel drill press on a workstation stand (Dremel 220-01 workstation) which allowed for stability, control, and precision during cutaneous wounding. The drill bit was a 1/16-inch micro grain carbide square end mill with 4 flutes from Speed Tiger Precision Technology (X002VBA0QV). The injuries in this study were performed with the aid of the Leica M60 stereomicroscope for increased magnification.

### Cutaneous wound preparation

Adult zebrafish from 3 to 13 months old were anesthetized in 84 mg/l Tricaine (0.5X Tricaine) in system water (Tricaine S by Syndel ver. 121718, buffered to pH7 with 1M Tris-pH9), and then moved to 168 mg/l Tricaine (1X Tricaine) in system water for brief final anesthetization before injury, as described in the results. Gradual anesthetization using Tricaine appears to be better for adult fish survival, especially for live imaging. To induce a cutaneous wound, fish were loaded onto the holding platform on top of a strip Whatman filter paper soaked in 1X Tricaine to help keep the fish moist and anesthetized. 3D-printed anchoring rods were then inserted into their holders to keep fish immobilized during injury, and mounted fish were moved to the stereomicroscope where the rotary tool was positioned next to them. After the cowling and drill bit were properly lined up below the midpoint of the fish between the pelvic and anal fin, the rotary tool was turned on at speed 3 (8500 RPM) and gently pushed into the left side of the fish for 10 seconds to create a small cutaneous wound. After injury fish were returned to system water and revived.

### BODIPY Staining

BODIPY dye was used to stain cell membranes in living zebrafish using a similar protocol as previously described (Lumaquin et al., 2021). BODIPY far-red fluorescent dye was reconstituted in 1ml of DMSO to make a 5mg/ml stock solution. Adult zebrafish were bathed in a 20ng/ml BODIPY bath (2ul of BODIPY Staining Stock Solution (ThermoFisher Scientific D10000) in 50 mL of system water) for at least 2 hours. After the bath fish were rinsed for one hour in clean system water before imaging.

### Image acquisition

Confocal images of skin re-epithelialization, neutrophil recruitment, and blood and lymphatic vessel regrowth were taken on the Nikon Ti2 inverted microscope with Yokogawa CSU-W1 spinning disk confocal, Hamamatsu Orca Flash 4 v3 camera with the following Nikon objectives: 2× Air 0.1 N.A., 4× Air 0.2 N.A., 10× Air 0.45 N.A. Expression of *Tg(actb2:GFP)* and Tg(mrc1a:eGFP)^y251^ was attained using a 488nm excitation laser. Tg(lyz:DsRed)^nz50^, Tg(kdrl:mcherry)^y206^ expression was acquired using a 561nm excitation laser, and BODIPY expression was taken using a 633nm excitation laser. The 405nm excitation laser was used to obtain autofluorescence in the scales, which allowed for easy visualization of the wound area. For single timepoint images fish were anesthetized and mounted into a chambered coverglass filled with 84-168mg/l of Tricaine water with the wound side facing down and covered with a moistened sponge to hold the fish in place. Overnight time-lapse images of skin re-epithelialization and neutrophil recruitment after wounding, Movie 2 and Movie 3 respectively, were acquired using an adult fish intubation protocol as previously described (Castranova et al., 2022). Stereomicroscope pictures of fish were taken using the Leica M165 FC microscope with a Leica DFC 7000 T camera. Pictures of the injury setup apparatus (Fig. 1) were taken using a Sony α 6400 mirrorless camera and iPhone 11. Video of the injury setup and generation of a cutaneous wound in zebrafish using a rotary tool (Movie 1) was filmed using an iPhone 11 and edited in iMovie and Adobe Premiere Pro 2022.

### Image processing and analysis

Confocal images were processed using Nikon Elements and maximum-intensity projections of confocal stacks are shown. Time-lapse movies of skin re-epithelialization (Movie 2) and neutrophil recruitment (Movie 3) were processed in Nikon Elements and transferred to Adobe Premiere Pro 2022 for compiling and editing. Schematics and figures were made using Adobe Photoshop 2022, Adobe Illustrator 2022, and BioRender software. Wound size area, blood and lymphatic vessel regrowth area, and neutrophil fluorescent intensity were measured using Fiji software. Neutrophil fluorescence was measured based on the sum intensity projections of 4x confocal stacks with a 12.5μm step size. The corrected total cell fluorescence (CTCF) was calculated based on the postinjury wound area for each timepoint, by taking the integrated density of the wound area and subtracting the wound area times the mean of 3 fluorescence background readings. The CTCF for each timepoint was normalized over the CTCF of the preinjury image from the corresponding fish which was calculated in the same manner.

### Statistical analysis

For all quantifications, *n* represents the number of fish analyzed. For neutrophil fluorescence measurements statistical significance was expressed as P values and determined using a ratio paired t test comparing the preinjury CTCF levels to the CTCF level of each fish at each timepoint. All statistical tests were run, and graphs generated using PRISM 9 software. Error bars indicate the mean and standard deviation (Fig. 2) or mean and standard error (Fig. 3-4). (*) denotes p<0.05, (**) denotes p<0.01.

### Zebrafish husbandry and fish strains

Fish were housed in a large zebrafish-dedicated recirculating aquaculture facility in 6 l and 1.8 l tanks. Water quality parameters were routinely measured, and appropriate measures were taken to maintain water quality stability (water quality data available upon request). Fry were fed rotifers and adults were fed Gemma Micro 300 (Skretting) once per day. The following transgenic fish lines were used for this study: *Tg(mrc1a: eGFP)^y251^* (Jung et al., 2017), *Tg(kdrl:mcherry)^y206^* (Gore et al., 2011), *Tg(lyz:DsRed2)^nz50^* (Hall et al., 2007), *Tg(actb2:GFP)*(Higashijima et al., 1997). Fish used to image neutrophil recruitment and vessel regrowth were maintained and imaged in a *casper* (*roy^a9^* (Ren et al., 2002) plus *nacre^w1^* (Lister et al., 1999)) double mutant (White et al., 2008) genetic background for better visualization in the absence of melanocyte and iridophore cell populations. This study was performed in an AAALAC accredited facility under an active research project overseen by the NICHD ACUC, Animal Study Proposal 21-015 for zebrafish.

## Acknowledgments

The authors thank members of the Weinstein laboratory for their helpful comments on this manuscript. We thank Dr. Ten-Tsao Wong for the *Tg(actb2:GFP)* fish and Erik Skantze for drill bit procurement. The authors also thank the Research Animal Branch of the Eunice Kennedy Shriver National Institute of Child Health and Human Development as well as the Charles River staff for excellent animal care and husbandry.

## Author Contributions

Conceptualization: L.J.G., D.C., B.M.W.; Methodology: L.J.G., K.A., D.C., B.M.W.; Validation: L.J.G., K.A., D.C; Formal analysis: L.J.G., K.A.; Investigation: L.J.G., K.A., D.C., B.M.W.; Resources: B.M.W.; Data curation: L.J.G., K.A.; Writing - original draft: L.J.G., K.A., B.M.W.; Writing - review & editing: L.J.G., K.A., D.C., B.M.W.; Visualization: L.J.G., K.A., B.M.W.; Supervision: L.J.G., B.M.W.; Project administration: B.M.W.; Funding acquisition: B.M.W

## Movies

**Movie 1: Cutaneous wounding in zebrafish.**

Movie showing preparation and generation of a cutaneous wound in zebrafish using a rotary tool. The following steps are shown: 1) addition of filter paper with Tricaine water to the holding platform, 2) loading of the fish on to the holding platform, 3) securing fish on to the holding platform using anchoring rods, 4) rotating insert to angle fish, 5) lining up the rotary tool to the site of injury, 6) creating the injury with the rotary tool, 7) final wounding outcome.

**Movie 2: Time-lapse imaging of skin re-epithelialization after wounding.**

Still and timelapse images of a *Tg(actb2:GFP)* (βactin) fish before and after cutaneous wounding. **0-5”** Still images of fish prior to injury including a zoomed-out image and close-up image of the wound site. **6-16”** Time-lapse of skin re-epithelization 0.5 – 17.6 hours post injury. **17-19”** High magnification time-lapse 18 hours post injury showing the infiltration of many cell types into the wound area. **20-24”** Still image of wound area 2 days post injury showing the wound has been closed.

**Movie 3: Neutrophil recruitment after cutaneous wounding.** Still and timelapse images of a Tg(lyz:dsRed) fish stained with BODIPY before and after cutaneous wounding. Schematic of fish shows approximate site of wounding. **0-4”** Still image of wound area before injury shows scales intact and neutrophils evenly scattered throughout. **5-16”** Time-lapse of neutrophil recruitment 0.5 – 17.25 hours post injury. A mass influx of neutrophils is seen during the indicated time suggesting that neutrophils respond quickly to cutaneous injury. BODIPY labeling shows skin re-epithelialization occurring simultaneously to neutrophile recruitment.

## Supplemental FIles

**Supp. File 1: CAD design for plastic rotary tool cowling.**

CAD design for custom 3D printing of a plastic cowling to fit over he end of the rotary tool, to limit bit depth during wounding.

**Supp. File 2: CAD design for plastic holding platform.**

CAD design for custom 3D printing of a plastic holding platform with a rotating insert, to immobilize fish and allow them to be positioned for the injury procedure.

**Supp. File 3: CAD design for plastic rod for holding platform.**

CAD design for custom 3D printing of a plastic rod used to pin a fish to the rotating insert in the holding platform, to immobilize the fish for the injury procedure.

**Supp. File 4: CAD design for plastic rotary tool cowling.**

CAD design for custom 3D printing of a plastic rod used to pin a fish to the rotating insert in the holding platform, to immobilize the fish for the injury procedure.

